# Surfactin accelerates *Bacillus subtilis* pellicle biofilm development

**DOI:** 10.1101/2024.10.13.618088

**Authors:** Rune Overlund Stannius, Sarah Fusco, Michael Cowled, Ákos T. Kovács

**Author notes:** Corresponding author: Ákos T. Kovács.

## Abstract

Surfactin is a biosurfactant produced by many *B. subtilis* strains with a wide variety of functions from lowering surface tension to allowing motility of bacterial swarms, acting as a signaling molecule, and even exhibiting antimicrobial activities. However, the impact of surfactin during biofilm formation has been debated with variable findings between studies depending on the experimental conditions.

*B. subtilis* is known to form biofilms at the solid-air, the solid-medium, and the liquid-air interfaces, the latter of which is known as a pellicle biofilm. Pellicle formation is a complex process requiring coordinated movement to the liquid-air interface and subsequent cooperative production of biofilm matrix components to allow robust pellicle biofilm formation. This makes pellicle formation a promising model system for assaying factors in biofilm formation and regulation.

Here, we assayed the influence of surfactin and additional metabolites on the timing of pellicle biofilm formation. Using of time-lapse imaging, we assayed pellicle formation timing in 12 *B. subtilis* isolates and found that one, MB9_B4, was significantly delayed in pellicle formation by approximately 10 hours. MB9_B4 was previously noted to lack robust surfactin production. Indeed, deletion of surfactin synthesis in the other isolates delayed pellicle formation. Further, pellicle delay was rescued by addition of exogeneous surfactin and spent media from mature pellicles. Testing reporters of biofilm-related gene expression revealed that induction of pellicle formation was caused by a combination of increased gene expression of one of the biofilm components and promotion of growth. Intriguingly, spent media of surfactin mutant strains were also able to stimulate pellicle formation, indicating possible additional metabolites also influence the timing of pellicle development.

## 1. Introduction

Biofilm is one of the most common lifestyles of bacteria in natural settings [1–4]. During biofilm formation, microorganisms produce a robust extracellular matrix consisting of polysaccharides, proteins, and eDNA, which together provide protection against antibiotics, predators, invasion or non-cooperative microorganisms, and other environmental factors [5–8]. Formation of biofilm is a complex coordinated effort in which spatial gene expression patterns [9,10], division of labor [11], and cell differentiation [12,13] are common, and these make biofilm development a highly relevant target within the field of sociomicrobiology [14,15]. Additionally, biofilm and the dynamics of their formation are highly relevant in our society as an agent of persistent bacterial colonization in sickness and in health of humans, animals, and plants, as well as industrial production and infrastructure [16].

Growing in a biofilm is a tradeoff with accompanying detrimental effects on the microbes within it; most biofilms are static limiting space and nutrients; potentially negatively affecting fitness if formed indiscriminately in an ever changing environment [17,18]. Therefore, biofilm development is tightly regulated and tuned including precise gene regulatory pathways being activated during establishment, maturation, and disassembly toward a search for a new niche to colonize [19]. Biofilms are generally studied using distinct laboratory models, including architecturally complex colony biofilms at the interface between a solid substrate and air, submerged biofilms are formed at the substrate-liquid interface, floating or host-embedded aggregates, and pellicle biofilms which are formed floating at the liquid-air interface [20]. These biofilm types have specific requirements to establish successfully, and therefore it is also important that the regulation of biofilm related processes either can accommodate several types of modalities or that biofilm itself is robustly functional in several situations.

Pellicle biofilms are in particular dependent on precise coordination to form a robust biofilm requiring coordinated movement and production of matrix to succeed [11,12]. Unlike colony biofilms, which can arise from clonal growth in one spot, cells during pellicle biofilm development transition from a planktonic phase in the liquid medium into a sessile state in the floating biofilm through a well-defined time window which leaves the lower liquid medium almost devoid of cells [14,21,22]. This behavior is easily observable and provides a good tractable model system for studying the factors determining biofilm formation and the coordination of differentiation.

*Bacillus subtilis* is a Gram positive, soil-dwelling bacterium with well-documented plant growth promoting abilities through direct inhibition of plant pathogens as well as through induced systemic resistance and promotion of plant growth [23]. Plant protection by *B. subtilis* is mediated by a range of bioactive molecules such as lipopeptides which have been shown as a direct effector of plant growth and resistance. One such molecule is surfactin, a surface active agent involved in surface colonization by *B. subtilis* [24–29] and abundantly produced during colonization of plant roots where it also induces systemic resistance [30–34]. Importantly, surfactin promotes *B. subtilis* to reach and colonize the rhizosphere when inoculated to the soil [35] and domesticated *B. subtilis* strains that lack surfactin production demonstrate reduced root colonization in the soil compared with undomesticated strains, unlike during direct inoculation of the strain on the seedlings in agar medium [36]. *B. subtilis* pellicle biofilm formation largely depends on two essential matrix components [37,38], exopolysaccharides (EPS) encoded by the *epsABCDEFGHIJKLMNO* operon (*epsA-O*) [39] and protein fibers of TasA, encoded by the *tapA-sipW-tasA* (*tapA*) operon [40].

While the importance of surfactin is clear in collective swarming motility and for its effects on plants, its impact on biofilm formation is controversial. A previous study found no significant difference in biofilm mass on tomato roots between wild-type strains and their surfactin biosynthetic gene cluster (BGC) deletion mutants of 5 *B. subtilis* root-derived isolates [41], others noted lack of biofilm formation and tomato colonization by the surfactin deletion mutant of *B. subtilis* 6051 [30]. Within the *B. subtilis* group species, *B. velezensis* (FZB42) and *B. amyloliquefaciens* (UMAF6614) were reported to have severe biofilm defects when surfactin biosynthesis operon was disrupted [42,43]. Indeed, disruption of surfactin biosynthesis was reported to have a species-dependent influence on pellicle formation in the *B. subtilis* group [44].

Previous reports demonstrated the direct influence of surfactin on induction of biofilm-related gene expression in the model strain *B. subtilis* NCIB3610 (hereafter 3610) when cells are grown in exponential growth phase; thus, practically non-biofilm inducing conditions [45–47]. Subsequent studies interpreted these results that surfactin is crucial for biofilm development of *B. subtilis*. Therefore, the essentiality of surfactin for biofilm formation has been previously revisited, which demonstrated that deletion of the surfactin biosynthetic gene *srfAA* in 3610 and six other recent isolates has no observable influence on pellicle formation, monitored after 20 h, and root colonization in biofilm inducing MSgg and MSNg media using hydroponic conditions [48]. Additionally, re-sequencing of the genome of the originally created and widely tested *srfAA* strain that displays reduced pellicle biofilm formation revealed additional point mutations, which likely explain disrupted biofilm development, especially, as re-introduction of the same *srfAA* mutation again into 3610 had no apparent influence on biofilm development after 20 h [48].

In this study, we dissect the influence of surfactin on biofilm development. When employing time lapse experiment, we observed a significant delay of pellicle formation for the soil isolate MB9_B4, a recent natural *B. subtilis* isolate that lacks robust surfactin production. The delayed pellicle development of MB9_B4 could be rescued by the addition of spent media supernatant from either the wild-type MB9_B4 or its surfactin deletion derivative. Our data demonstrates that surfactin promotes an early development of pellicles in *B. subtilis*, in addition to a distinct yet to be identified factor, validating the influence of surfactin on biofilm development, while not being essential for its pellicle formation.

## 2. Materials and methods

### 2.1 Strains, chemicals, and genetic modification

Strains used in this study can be found in **Table 1** and have earlier been genomically and chemically characterized in [49]. Strains were routinely cultured in lysogeny broth (LB; Lennox, Carl Roth, 10 g/L tryptone, 5 g/L yeast extract and 5 g/L NaCl) and LBgm (LB supplemented with 1% v/v glycerol and 0.1 mmol/L MnCl_2_, based on [50]). Antibiotics were used at the following final concentrations: Spectinomycin (spec) 100 µg/mL, kanamycin (kan) 5 µg/mL, tetracycline (tet) 10 µg/mL, chloramphenicol (chl) 10 µg/mL.

All strains were naturally competent and genetically engineered using a modified version of the transformation protocol described in [51]. Briefly, 1 mL of overnight culture was spun down and resuspended in 100 µL sterile MQ water of which 10 µL were inoculated into 2 mL competence medium (80 mmol/L K_2_HPO_4_, 38.2 mmol/L KH_2_PO_4_, 20 g/L glucose, 3 mmol/L Na_3_-citrate, 45 µmol/L ferric NH_4_-citrate, 1 g/L casein hydrolysate, 2 g/L K-glutamate, 0.335 µmol/L MgSO_4_·7H_2_O), and incubated at 37 °C for 3.5 hours. Donor DNA was extracted from 1 mL of overnight grown culture using the Bacterial and Yeast Genomic DNA Purification Kit from EURx with a typical purified DNA concentration ranging from 50 – 150 ng/µL. 2 µl donor DNA was added to a new tube and washed down using 400 µL of competent cells and incubated for further 2 hours before plating 100 µL on selective LB agar plates at 37 °C overnight to select for successful transformants.

To construct DK1042 P*_srfAA_*-*gfp*, the P*rrnB* was replaced with the *srfAA* promoter in the pGFP-rrnB vector [52] using prolonged overlap extension PCR. The *srfAA* promoter region was PCR-amplified with the primers srfAA_forward (5’ AGCTGTCAAACATGAGAATTGAAA GAATCGTTGTAAGACGC 3’) and srfAA_reverse (5’ AGTTCTTCTCCTTTGCTAGCTTATTTCCATATTGTC ATACCTCC 3’). pGFP-rrnB was linearized via PCR with pGFP_forward (5’ GTATGACAATATGGAAA TAAGCTAGCAAAGGAGAAGAACT 3’) and pGFP_reverse (5’ CGTCTTACAACGATTCTTTCAATTCTCAT GTTTGACAGCTT 3’) using Q5 DNA polymerase. The PCR products were directly transformed into *Escherichia coli* cells, and correct insert was verified by sequencing. The construct was introduced into *B. subtilis* DK1042 *amyE* locus using natural competence selecting for chloramphenicol resistance and verified by sequencing.

**Table S1:**
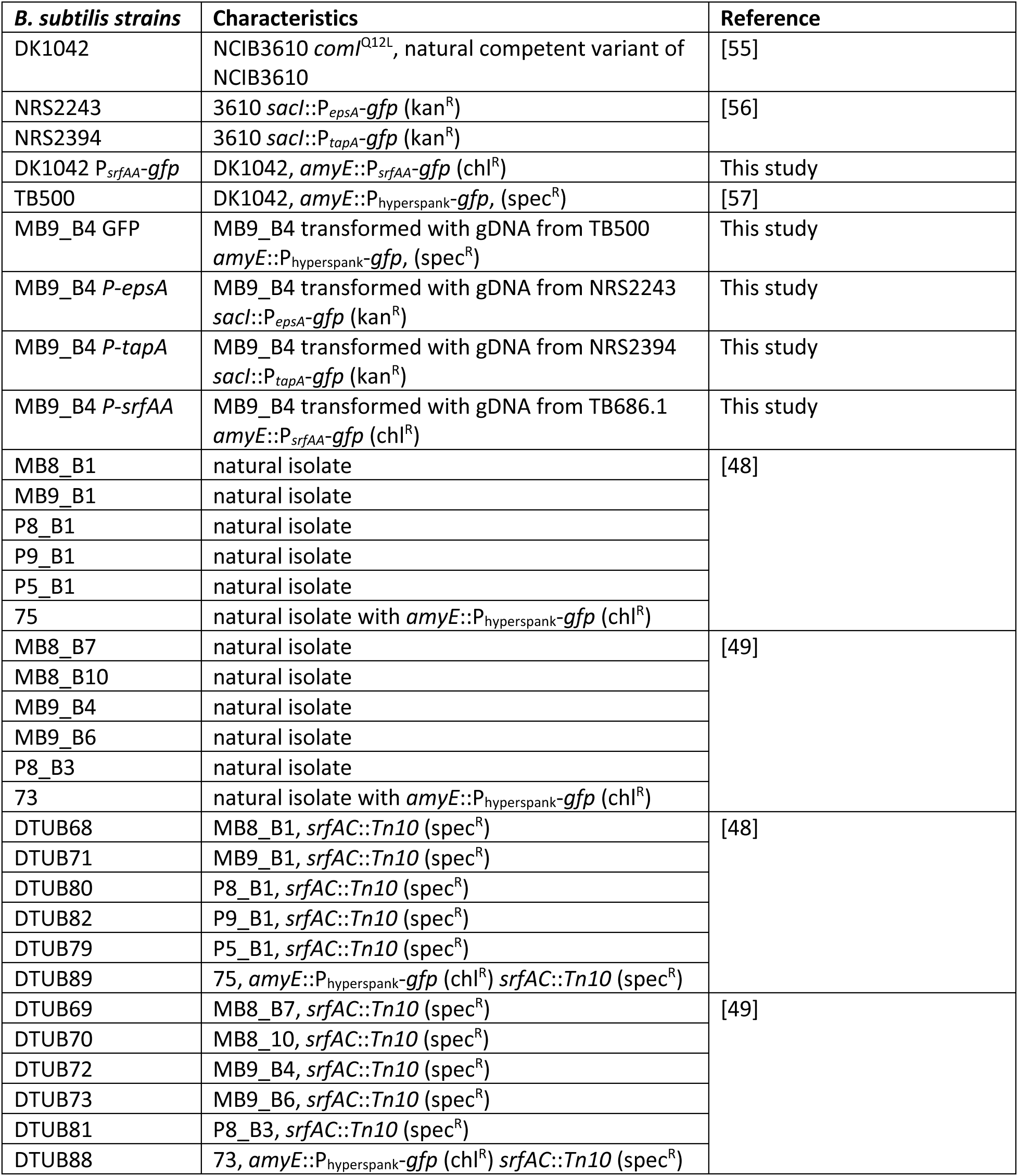
strains used in the study. *gfp: gfpmut2*, kan^R^: kanamycin resistance, chl^R^: chloramphenicol resistance, spec^R^: spectinomycin resistance

| ***B. subtilis* strains** | **Characteristics** | **Reference** |
| --- | --- | --- |
| DK1042 | NCIB3610 *comI*<sup>Q12L</sup>, natural competent variant of NCIB3610 | [55] |
| NRS2243 | 3610 *sacI*::*P<sub>epsA</sub>-gfp* (kan<sup>R</sup>) | [56] |
| NRS2394 | 3610 *sacI*::*P<sub>tapA</sub>-gfp* (kan<sup>R</sup>) |  |
| DK1042 *P<sub>srfAA</sub>-gfp* | DK1042, *amyE*::*P<sub>srfAA</sub>-gfp* (chl<sup>R</sup>) | This study |
| TB500 | DK1042, *amyE*::*P<sub>hyperspank</sub>-gfp*, (spec<sup>R</sup>) | [57] |
| MB9_B4 GFP | MB9_B4 transformed with gDNA from TB500 *amyE*::*P<sub>hyperspank</sub>-gfp*, (spec<sup>R</sup>) | This study |
| MB9_B4 *P-epsA* | MB9_B4 transformed with gDNA from NRS2243 *sacI*::*P<sub>epsA</sub>-gfp* (kan<sup>R</sup>) | This study |
| MB9_B4 *P-tapA* | MB9_B4 transformed with gDNA from NRS2394 *sacI*::*P<sub>tapA</sub>-gfp* (kan<sup>R</sup>) | This study |
| MB9_B4 *P-srfAA* | MB9_B4 transformed with gDNA from TB686.1 *amyE*::*P<sub>srfAA</sub>-gfp* (chl<sup>R</sup>) | This study |
| MB8_B1 | natural isolate | [48] |
| MB9_B1 | natural isolate |  |
| P8_B1 | natural isolate |  |
| P9_B1 | natural isolate |  |
| P5_B1 | natural isolate |  |
| 75 | natural isolate with *amyE*::*P<sub>hyperspank</sub>-gfp* (chl<sup>R</sup>) |  |
| MB8_B7 | natural isolate | [49] |
| MB8_B10 | natural isolate |  |
| MB9_B4 | natural isolate |  |
| MB9_B6 | natural isolate |  |
| P8_B3 | natural isolate |  |
| 73 | natural isolate with *amyE*::*P<sub>hyperspank</sub>-gfp* (chl<sup>R</sup>) |  |
| DTUB68 | MB8_B1, *srfAC*::*Tn10* (spec<sup>R</sup>) | [48] |
| DTUB71 | MB9_B1, *srfAC*::*Tn10* (spec<sup>R</sup>) |  |
| DTUB80 | P8_B1, *srfAC*::*Tn10* (spec<sup>R</sup>) |  |
| DTUB82 | P9_B1, *srfAC*::*Tn10* (spec<sup>R</sup>) |  |
| DTUB79 | P5_B1, *srfAC*::*Tn10* (spec<sup>R</sup>) |  |
| DTUB89 | 75, *amyE*::*P<sub>hyperspank</sub>-gfp* (chl<sup>R</sup>) *srfAC*::*Tn10* (spec<sup>R</sup>) |  |
| DTUB69 | MB8_B7, *srfAC*::*Tn10* (spec<sup>R</sup>) | [49] |
| DTUB70 | MB8_B10, *srfAC*::*Tn10* (spec<sup>R</sup>) |  |
| DTUB72 | MB9_B4, *srfAC*::*Tn10* (spec<sup>R</sup>) |  |
| DTUB73 | MB9_B6, *srfAC*::*Tn10* (spec<sup>R</sup>) |  |
| DTUB81 | P8_B3, *srfAC*::*Tn10* (spec<sup>R</sup>) |  |
| DTUB88 | 73, *amyE*::*P<sub>hyperspank</sub>-gfp* (chl<sup>R</sup>) *srfAC*::*Tn10* (spec<sup>R</sup>) |  |

### 2.2 Culture conditions

For pellicle timing assays, overnight incubated cultures in LB medium were adjusted to 0.1 at optical density at 600 nm (OD_600_) using LB medium and 10 µl were inoculated into 1 mL of LBgm in 24-well microtiter plates and were incubated at 30 °C and followed using the ReShape imaging system (ReShape biotech, Denmark) which allowed time-lapse imaging of each microtiter well every 30 minutes over a period of 48 hours. When testing for induction, 10 µL of either the spent media or fresh LBgm was added while derivative fractions of spent media were solubilized in H_2_O 1% v/v DMSO and compared to pure H_2_O 1% v/v DMSO acting as control. We observed that well opacity was a possible marker for pellicle formation due to the low cell density in static cultures before assembly at the liquid-air interface. To set a definition for when a pellicle is formed, we manually annotated pellicle timing and cross-referenced with image analysis data from the ReShape interface to find that pellicle formation coincided with the average RGB red value reaching 100 (**Fig. S1**) which was used for determining pellicle formation timing in the rest of the study.

### 2.3 Promoter fusions

To assay potential induction by spent media and surfactin, overnight cultures were adjusted to 0.01 at OD_600_ and 10 µL was inoculated into 188 µL media in a 96-well microtiter plate, and 2 µL methanol-dissolved surfactin (final concentration of 20 µmol/L in the medium) or filter sterilized spent media was added with fresh LBgm, or methanol as controls. Growth and promoter fusion expression was followed by microplate reader (BioTek Synergy HTX Multi-mode Microplate Reader). Plates were incubated at 30 °C with continuous shaking and the following were measured every 10 minutes for 48h: OD_600_, green fluorescence for promoter-GFP fusion, red fluorescence for constitutively expressed mKate2 (Optics position: Bottom, GFP: ex: 485/20 em: 528/20, Gain: 60, mKate2: Ex: 590/20, Em: 635/20, Gain 60).

### 2.4 Fractionation

The filtered spent media (2L) was partitioned against ethyl acetate (1% v/v HCOOH) (2 × 2L), and the ethyl acetate was concentrated *in vacuo* to an oily residue (1.02 g). Chromatographic fractionation was carried out on a Biotage Isolera One instrument, by first dissolving the crude organic extract in 40% v/v MeCN/H_2_O (0.01% v/v HCOOH, 7mL) and wet-loaded onto a C_18_ Biotage column (50 g) equilibrated with 40% v/v MeCN/H_2_O (0.01% v/v HCOOH). The column was then eluted stepwise between 40-100% v/v MeCN/H_2_O (0.01% v/v HCOOH), with 5% incremental increases (150 mL each), resulting in 30 fractions (80 mL each).

### 2.5 Data-Dependent LC-ESI-HRMS/MS Analysis

Ultra-high-performance liquid chromatography–diode array detection–quadrupole time-of-flight mass spectrometry (UHPLC–DAD–QTOFMS) was performed on an Agilent Infinity 1290 UHPLC system (Agilent Technologies, Santa Clara, CA, USA) equipped with a diode array detector. Separation was achieved on a 150 × 2.1 mm i.d., 1.9 μm, Poroshell 120 Phenyl Hexyl column (Agilent Technologies, Santa Clara, CA) held at 40°C. The sample, 1 μL, was eluted at a flow rate of 0.35 mL/min using a linear gradient from 10% v/v acetonitrile (LC-MS grade) in Milli-Q water buffered with 20 mM formic acid increasing to 100% v/v in 10 min, staying there for 2 min before returning to 10% v/v in 0.1 min. Starting conditions were held for 3 min before the following run.

Mass spectrometry (MS) detection was performed on an Agilent 6545 QTOF MS equipped with Agilent Dual Jet Stream electrospray ion source (ESI) with a drying gas temperature of 250°C, a gas flow of 8 L/min, sheath gas temperature of 300°C and flow of 12 L/min. Capillary voltage was set to 4000 V and nozzle voltage to 500 V in positive mode. MS spectra were recorded as centroid data, at an *m/z* of 100–1700, and auto MS/HRMS fragmentation was performed at three collision energies (10, 20, and 40 eV), on the three most intense precursor peaks per cycle. The acquisition rate was 10 spectra/s. Data was handled using Agilent MassHunter Qualitative Analysis software (Agilent Technologies, Santa Clara, CA). Lock mass solution in 70 % v/v MeOH in water was infused in the second sprayer using an extra LC pump at a flow of 15 μL/min using a 1:100 splitter. The solution contained 1 μmol/L tributylamine (Sigma-Aldrich) and 10 μmol/L Hexakis (2, 2, 3, 3-tetrafluoropropoxy) phosphazene (Apollo Scientific Ltd., Cheshire, UK) as lock masses. The [M + H]^+^ ions (*m/z* 186.2216 and 922.0098, respectively) of both compounds were used.

## 3. Results

### 3.1 Soil isolates feature remarkably similar windows of pellicle formation timing and show variable dependence on surfactin

While previous testing of *B. subtilis* isolates revealed comparable biofilm development after 2 days of incubation [48], time lapse imaging exposed that the soil isolate MB9_B4 displayed a delayed pellicle formation by several hours compared with the closely related co-isolate MB9_B1. Despite delayed pellicle formation, the mature pellicle of MB9_B4 did not show any signs of decreased robustness, therefore allowing a comparative study of the factors influencing the timing of pellicle biofilm formation. Further time-lapse imaging of various *B. subtilis* isolates showed that all investigated strains featured similar timing of the first layer of the pellicle (20**±**1.45 hours) except MB9_B4 in which the first layer was first visible at 31 **±** 1.72 hours (**Fig. 1**). Previous characterization of the isolates demonstrated differential ability in lipopeptide production, with MB9_B4 lacking robust surfactin production compared with the other isolates [49]. We found a significant increase in pellicle formation timing for 9 of the 12 Δ*srfAC* derivatives, strains lacking surfactin synthesis, compared to the timing of their respective wild-type ancestor indicating a role of surfactin in timing of pellicle formation in some strains (**Table S1**). MB8_B10, MB9_B4, and 73 did not exhibit a significant difference between wild-type and Δ*srfAC* mutant, which for MB9_B4 might be explained by the isolate already lacking surfactin production, however, this is not the case for MB8_B10 or 73; thus, there may be other yet unknown factors that influence the timing of in pellicle formation.

**Fig. 1.**
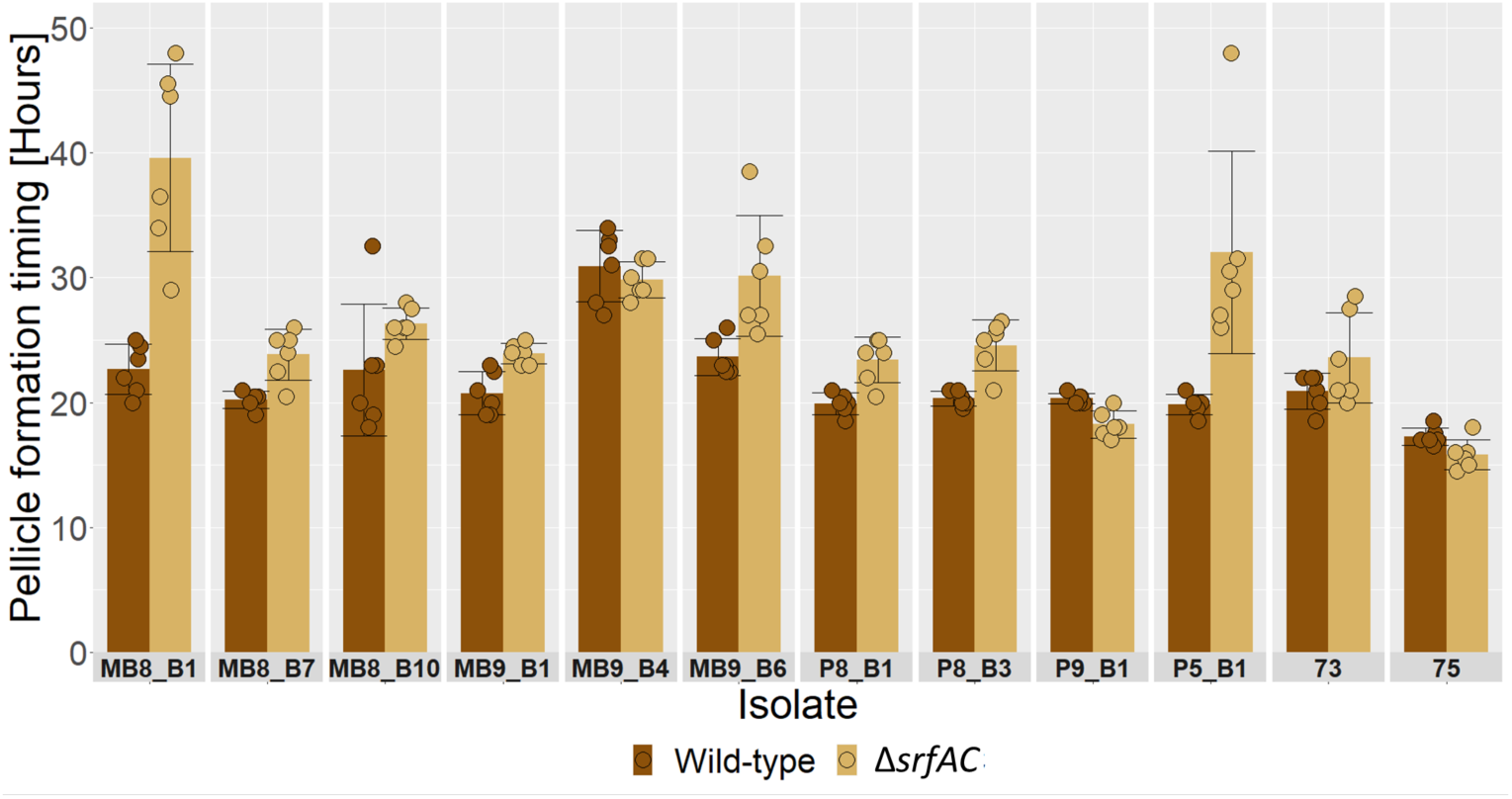
Average pellicle formation timing in LBgm at 30 °C for wild-type (dark brown) and Δ*srfAC* (beige) strains in rich biofilm inducing media with facets for each *B. subtilis* isolate separately. Average of 6 biological replicates, error bars represent standard deviation.

### 3.2 Surfactin and additional putative molecule(s) are able to rescue pellicle formation timing

As MB9_B4 is lacking robust surfactin production [49] and surfactin has been previously reported to induce expression of genes related to biofilm matrix production in exponential cell cultures [45], we wanted to test if complementation was possible by supplementation of pure surfactin or addition of sterilized spent media from mature pellicles (i.e. 48 h incubated cultures) from the collection of 12 soil isolates and their Δ*srfAC* derivatives.

Our results show that addition of 20 µM final concentration of surfactin was sufficient to ameliorate pellicle formation delay in MB9_B4 and furthermore significantly (**Table S2**) reduced pellicle formation timing in all but three of the tested Δ*srfAC* strains compared to that of the wild-type strain (**Fig. 3**).

**Fig. 2.**
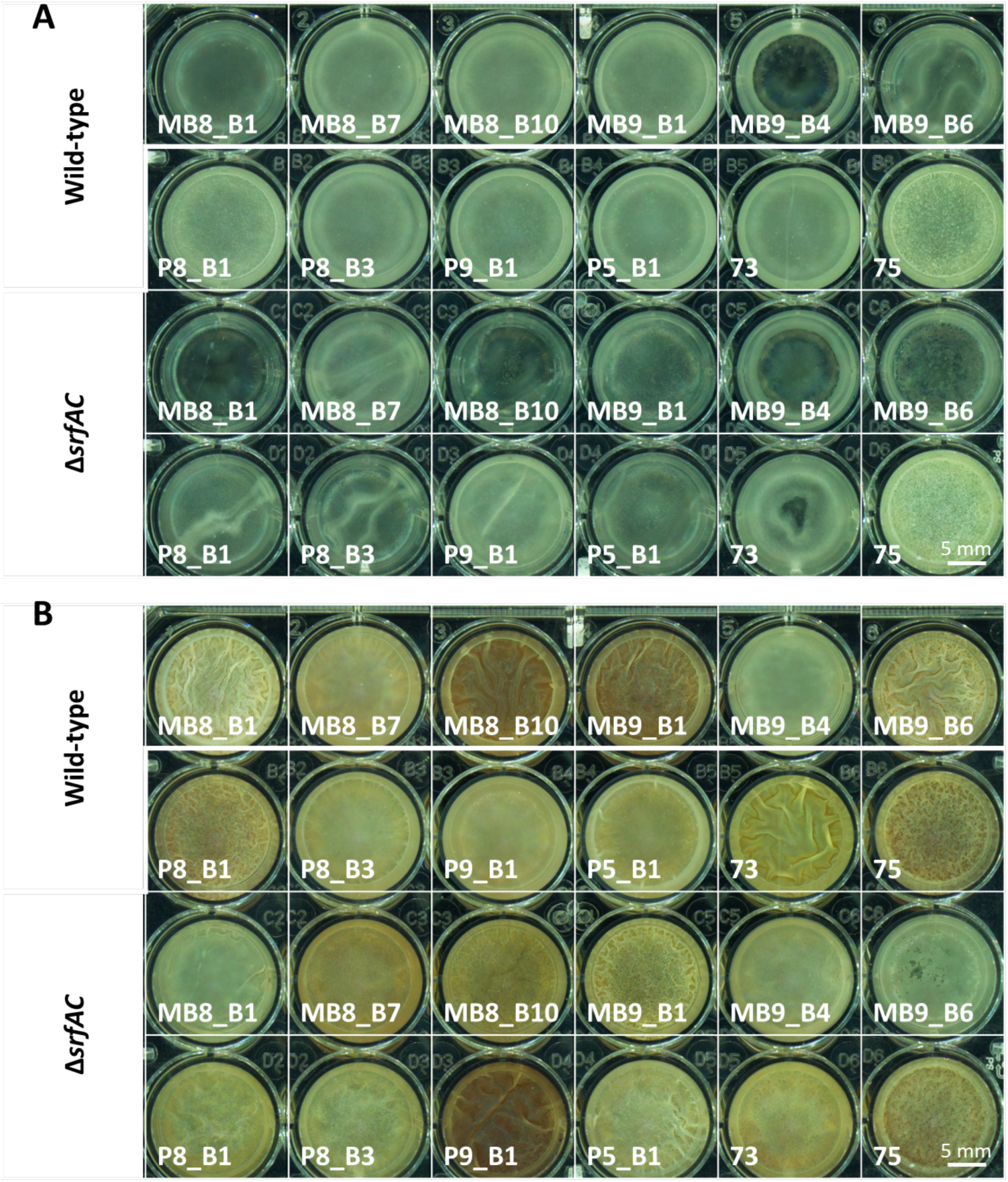
Pellicle biofilms cultured in 24-well plate at 30 °C, well diameter = 16.6 mm (5 mm scalebar shown in lower right corner of each 24-well plate). (A) Representative image of pellicle cultures at 20 hours showing early pellicles for most wild-type strains while the majority of Δ*srfAC* strains have not yet formed a pellicle. (B) Representative image of mature pellicle morphology at 48 hours illustrating robust pellicle formation by all tested strains regardless of surfactin production.

**Fig. 3.**
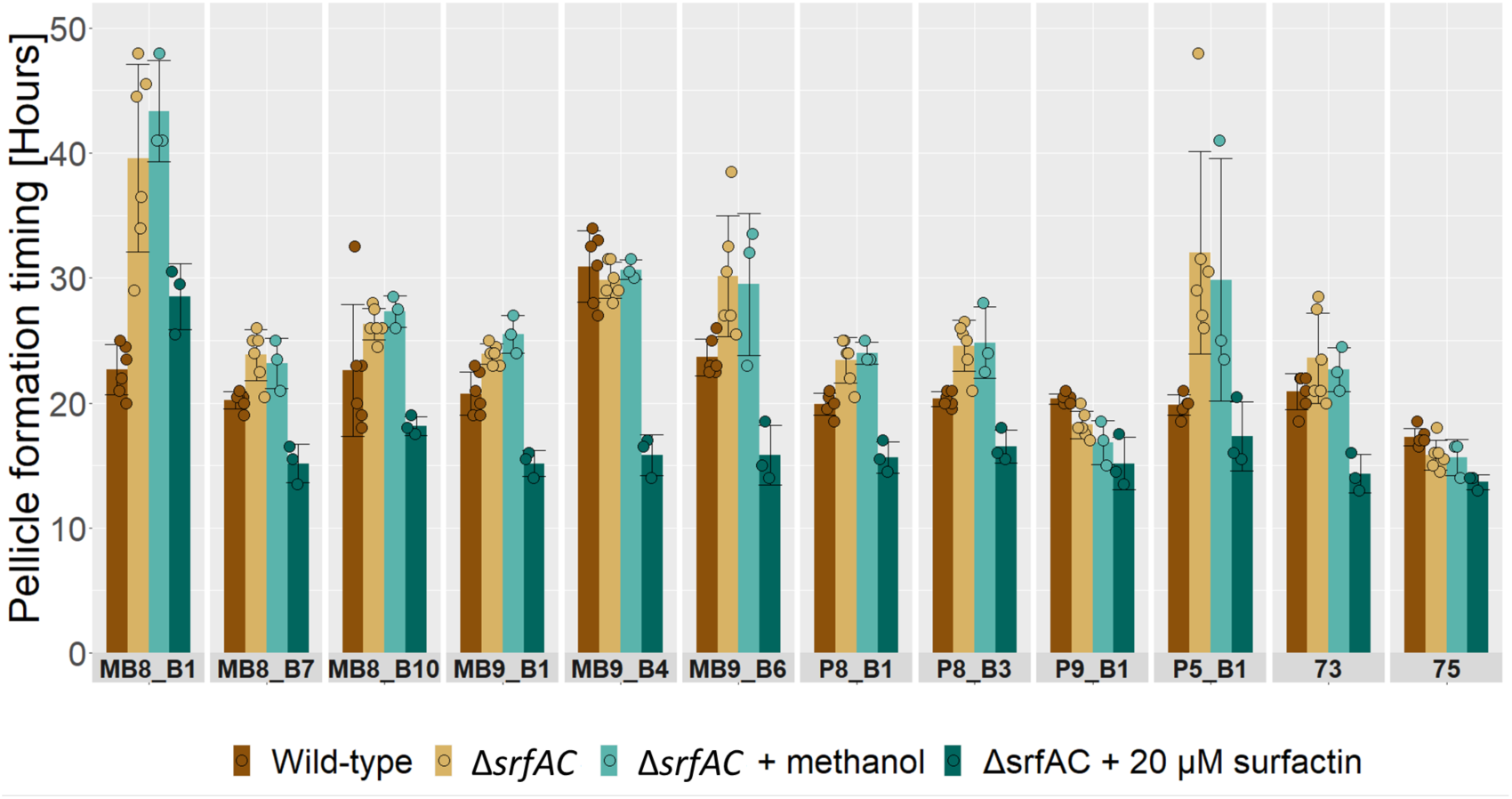
Induction of pellicle formation timing by addition of surfactin. Barplot showing average pellicle formation timing in LBgm at 30 °C for wild-type (dark brown), Δ*srfAC* (beige) for reference and Δ*srfAC* treated with methanol (light teal) or 20 µM final concentration of surfactin (dark teal) with facets for each *B. subtilis* isolate separately. Average of 6 biological replicates for wild-type and Δ*srfAC* and 3 biological replicates treated with methanol or surfactin, error bars represent standard deviation.

Moreover, addition of 0.1% v/v of spent media from mature pellicles of either wild-type and Δ*srfAC* versions of our strain collection was similarly able to hasten pellicle formation timing (**Fig. 4)**. MB8_B7, P8_B1, P8_B3, and P9_B1 inhibited early growth of MB9_B4 but still featured a reduction in pellicle formation time (**Fig. S2**). Earlier work has demonstrated that MB8_B7 contains the SPβ prophage encoding the bacteriocin sublancin which is present in spent media of the strain and can inhibit SPβ non-carriers [53]. Thus the observed inhibition could be an effect of kin discrimination factors against MB9_B4 as both P8_B1 and P8_B3 also carry the sublancin encoding SPβ prophage and P9_B1 carries the sporulation killing factor [49]. Overall, this suggested that lack of robust surfactin production in MB9_B4 might be responsible for the delayed biofilm development phenotype, as exogeneous surfactin could complement the deficiency. However, as indicated by complementation of spent media from Δ*srfAC* strains, there might be other unknown inducing compounds.

**Fig. 4.**
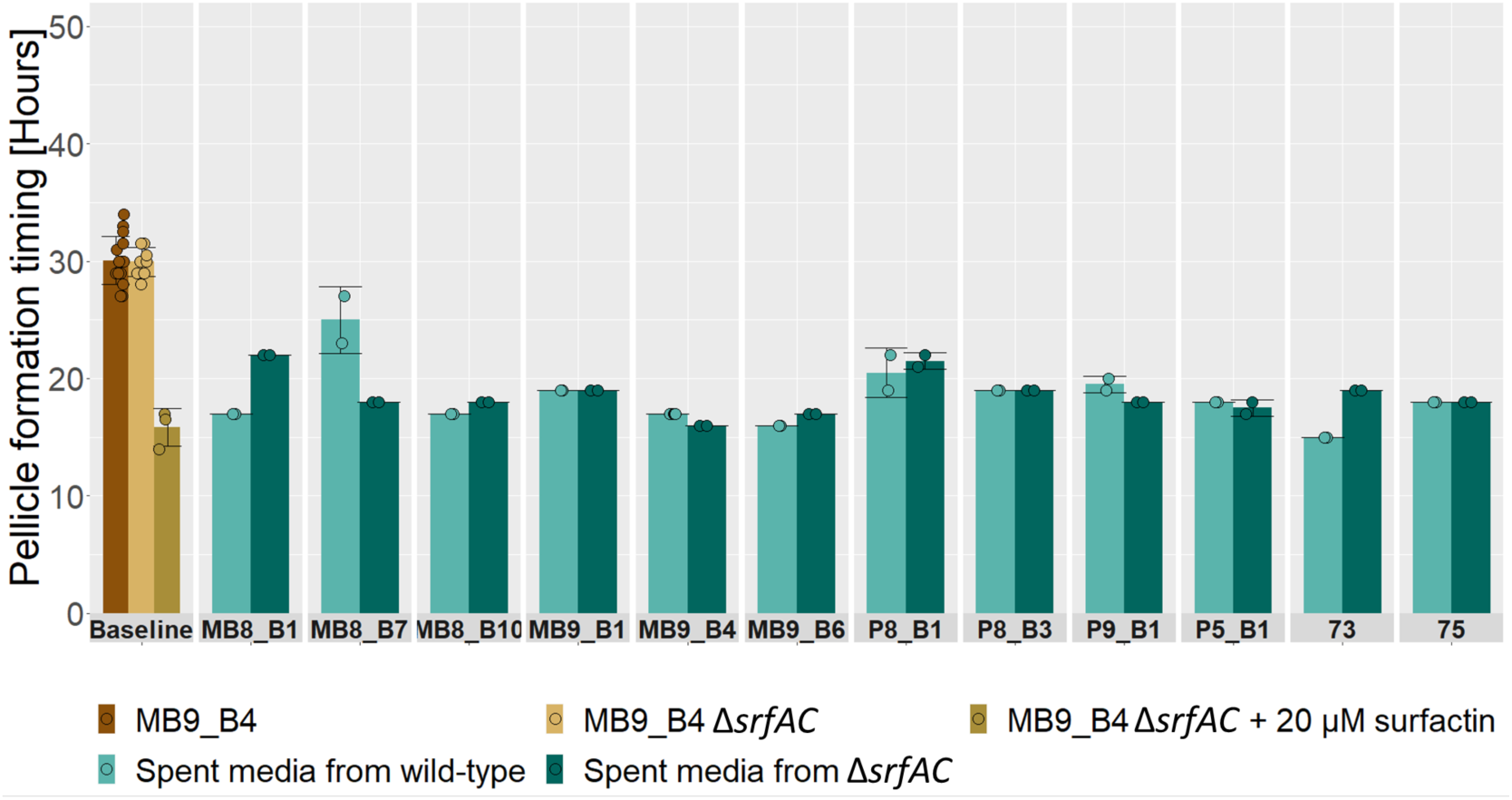
Induction of MB9_B4 pellicle formation by spent media from *B. subtilis* isolates and their Δ*srfAC* derivatives. Barplot showing average pellicle formation timing in LBgm at 30 °C for MB9_B4 wild-type (brown), MB9_B4 Δ*srfAC* (beige), and MB9_B4 Δ*srfAC* treated with 20 µM final concentration of surfactin (green-brown) in addition to MB9_B4 treated with spent media from either wild-type (light teal) or Δ*srfAC* (dark teal) with facets for each *B. subtilis* isolate separately. Average for baseline based on 8 biological replicates while spent media treatment is based on biological duplicates; error bars represent standard deviation.

To understand the mechanism by which surfactin is able to induce pellicle formation, we next assayed the effect of surfactin on the expression of biofilm matrix-related genes, *epsA* and *tapA* as well as on *srfA* itself using promoter fusions expressing GFP with an accompanying RFP tag under control of a constitutive promoter. Here, we observed little effect of surfactin on growth when judging by optical density at 600 nm (**Fig. 5A**), however, the constitutively active RFP tag seemed to exhibit increased fluorescence with addition of surfactin compared to the control (**Fig. 5B**), which could indicate increased growth. We do have to note that the RFP signal did not follow OD600 in early growth phases and might be an unreliable proxy for cell density, likewise, OD600 does not correlate well with cell density at higher densities which is where increase red fluorescence was observed. Additionally, an increase in green fluorescence from the reporter fusion which seemed independent from cell growth was observed for the *epsA* reporter fusion specifically with the addition of surfactin (**Fig. 5C**).

**Fig. 5.**
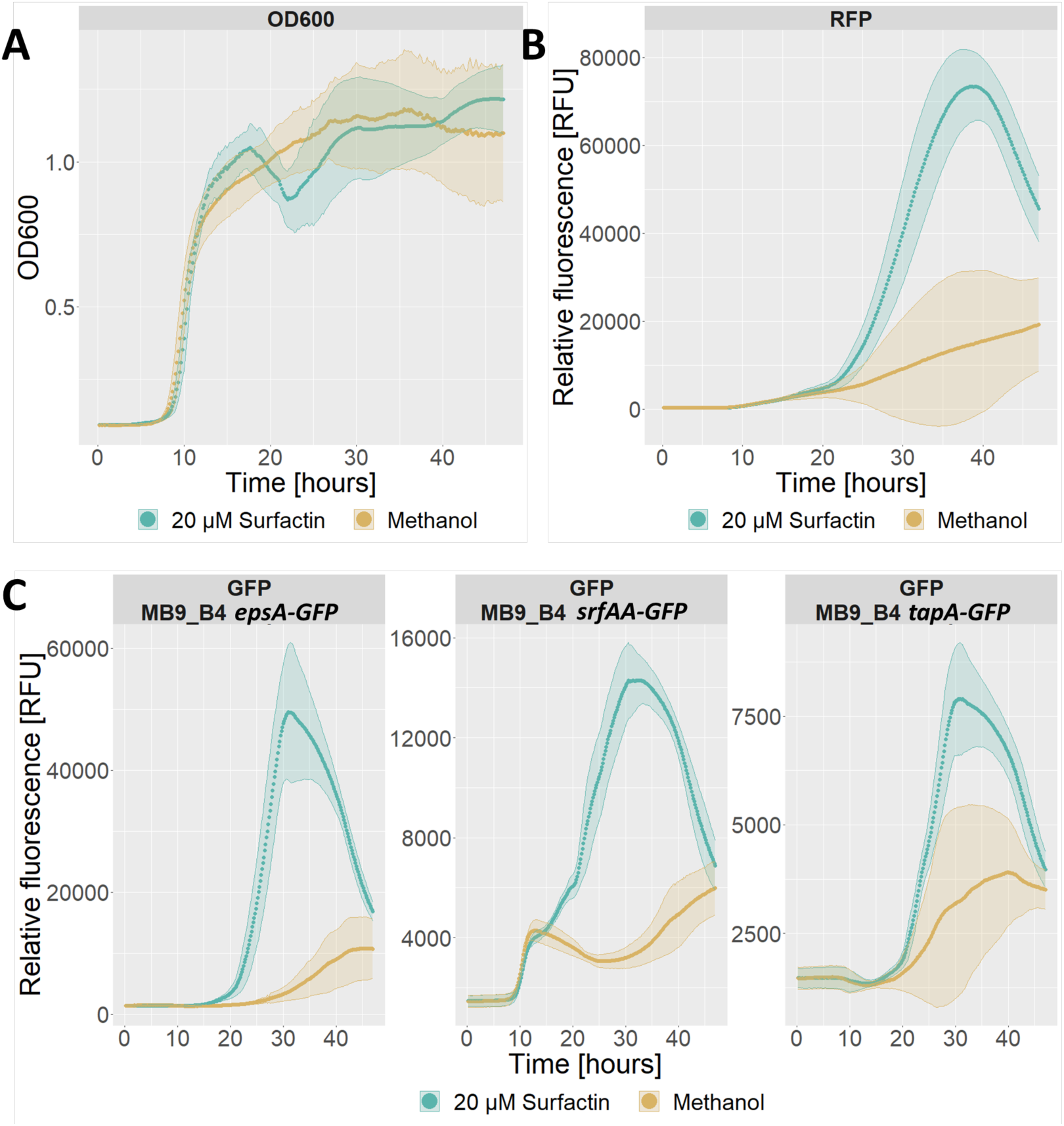
Effect of addition of surfactin to a final concentration of 20 µM (teal) or pure methanol as a control (beige) on growth and expression of fluorescence under constitutive and promoter-fusion coupled control in MB9_B4 over the course of 48 hours in LBgm at 30 °C. A) Optical density at 600 nm. B) Red fluorescence from RFP under control of a constitutive promoter. C) Green fluorescence from GFP under control of promoter fusions for *epsA* (left), *srfAA* (middle), or *tapA* (right). Data from all three strains pooled for A) and B), while C) is separated for each promoter fusion (split figures can be found in Fig. S3). Data averaged from 3 biological replicates with ribbons showing standard deviation.

To uncouple growth from gene expression, we adjusted GFP signal to account for live cell density by dividing the GFP signal value with either OD600 or the RFP signal. OD600 adjusted GFP signal showed an increase in expression of all three promoter fusions with the highest expression observed for *epsA* followed by *srfAA* and finally *tapA*, for all three promoter fusions, the increase in expression peaked around 30 hours and decreased towards the end of the assay (**Fig. 6A**). The RFP adjusted GFP expression on the other hand, only displayed a short window of increased expression for *epsA* from 20 to 30 hours and a reduction below the level of the control for *srfAA* from 25 hours and on (**Fig. 6B**). Overall, these results indicate that surfactin might be affecting expression of some biofilm components, primarily *epsA* in the later stages of growth in biofilm inducing media.

**Fig. 6.**
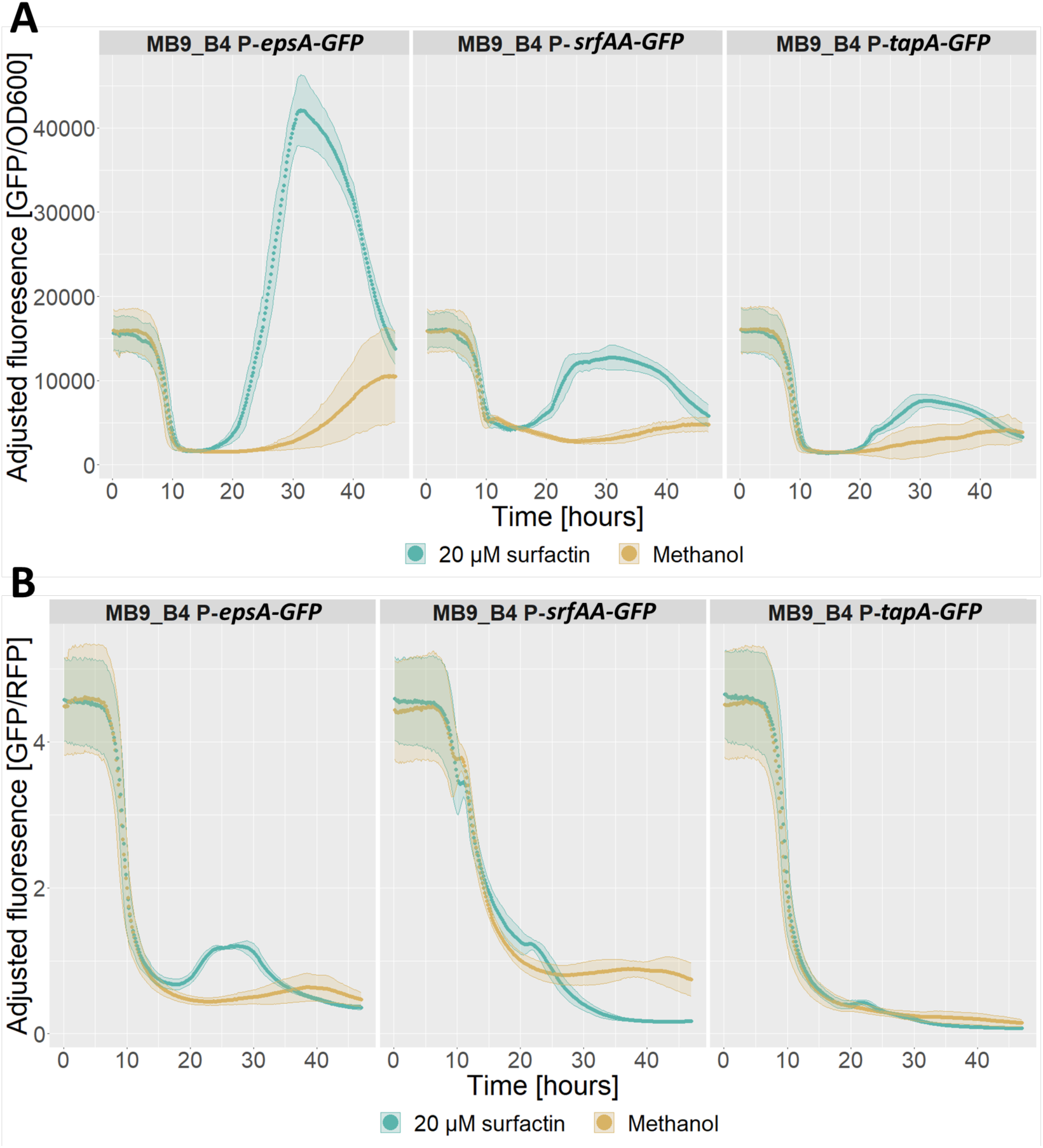
Adjusted reporter fusion expression of biofilm matrix-related genes *epsA* (left)*, tapA* (right), and surfactin biosynthesis connected *srfAA* gene (middle) in LBgm at 30 °C over the course of 48 hours with either pure methanol (beige) or treatment with surfactin to a final concentration of 20 µM (teal) A) GFP fluorescence signal adjusted by optical density at 600 nm. B) GFP fluorescence signal adjusted by red fluorescence. Data is the average of 3 biological replicates with ribbons showing standard deviation.

To identify other potentially inducing compounds, Solid Phase Extraction (SPE) was performed on the cell free supernatant of the biofilm model *B. subtilis* strain NCIB 3610, resulting in 30 fractions which were tested in the pellicle formation assay. Of the tested fractions, 7 were found to be advancing pellicle timing down to 15 hours, and 20 other fractions still featured reduced timing compared to the control. LC-ESI-HRMS/MS analysis of these fractions, however, only found surfactin to be specifically present in inducing fractions, and thus potential other inducing compounds are likely either lost during fractionation or eluted across several fractions at different concentrations complicating identification by presence/absence of induction.

## 4. Discussion

The role of surfactin in regulation of biofilm development of the *B. subtilis* species group has been tested in different laboratories [44,45,48]. Previous testing of 7 isolates and their derivatives lacking surfactin production for the ability to form pellicles and colonize plant root in liquid medium demonstrated that surfactin is not essential for either behavior in MSgg, MSNc pectin, and MOLP media, and that requirement of surfactin for biofilm development is likely species and media dependent [45,48]. Therefore, the role of surfactin as a universal signaling molecule required for pellicle biofilm development was questioned.

Revisiting the role of surfactin in induction of biofilm development, we sought to address the wide time-range at which pellicle formation is recorded. While some studies observed the presence of biofilms already after 8 hours, other studies recorded pellicle formation after 48 hours [44,45]; therefore, we employed time-lapse imaging to closely assay differences in biofilm development and induction of pellicle formation without a priori defining a time point when a pellicle should have been formed. Due to our temporal resolution, we were able to precisely follow differences in pellicle formation of 12 *B. subtilis* soil isolates and their derivative Δ*srfAC* mutants, lacking surfactin production. We observed that all tested isolates were able to produce robust pellicles; however, one isolate, MB9_B4, which lacks robust surfactin production displayed delay in pellicle formation production in rich biofilm inducing media. Furthermore, 9 out of 12 isolates were similarly delayed when their surfactin production capabilities were disrupted but were rescued by addition of exogeneous surfactin which decreased pellicle formation time sometimes below that of the wild-type for most strains.

We observed some indication that surfactin might increase growth in late stage culturing as evidenced by the increase in RFP expression under constitutive promoter control beyond 25 hours in our strains when treated with surfactin. However, we also have to note that RFP signal did not correspond well with OD600 in early growth where optical density still behaves linearly, and thus expression of RFP might have been affected in other ways than through increased growth alone. Promoter fusions reporting expression of biofilm related genes and the *srfA* operon, revealed an effect of surfactin on the expression of all three assayed genes with the most prominent effect observed for the *epsA* biofilm gene after signal adjustment with OD600 or RFP. Further experiments employing flow cytometry or single cell microscopy approaches are needed to uncouple expression levels from the cell density and allow a better understanding of the population dynamics of this induction – whether the whole population features increased expression or only a subset.

The growth promoting effect of surfactin could be due to increased oxygen diffusion, which surfactin has been shown to aid in earlier studies [54]. Pellicle formation dependence on cell density has been documented in earlier literature where other deficiencies could be complemented by higher starting cell density [12], which indicated that enhancement of cell division might be the possible mechanism for the observed advancement of pellicle timing.

Additionally, the ability of spent media of the Δ*srfAC* strains to speed up pellicle timing in MB9_B4 indicates that there may be other inducing compounds. Further mutational analysis or different fractionation strategies will be required to fully identify and confirm such putative biofilm development-promoting compound(s).

## Acknowledgments

This project was supported by the Danish National Research Foundation (DNRF137) for the Center for Microbial Secondary Metabolites and the Novo Nordisk Foundation for the “Imaging microbial language in biocontrol (IMLiB)” infrastructure grant (NNF19OC0055625).

## Supplementary information

**Table S1.** Bonferroni adjusted P-values of pairwise t-tests between wild-type and Δ*srfAC* mutant for each strain, * = below significance cut-off 0.05, NS = not significant.

| Isolate | Bonferroni corrected P-value | Isolate | Bonferroni corrected P-value |
| --- | --- | --- | --- |
| MB8_B1 | 3.34E-04 * | P8_B1 | 1.57E-03 * |
| MB8_B7 | 2.08E-03 * | P8_B3 | 6.20E-04 * |
| MB8_B10 | 1.21E-01 **NS** | P9_B1 | 1.32E-03 * |
| MB9_B1 | 2.22E-03 * | P5_B1 | 4.40E-03 * |
| MB9_B4 | 4.23E-01 **NS** | 73 | 1.25E-01 **NS** |
| MB9_B6 | 1.03E-02 * | 75 | 3.20E-02 * |

**Table S2.**
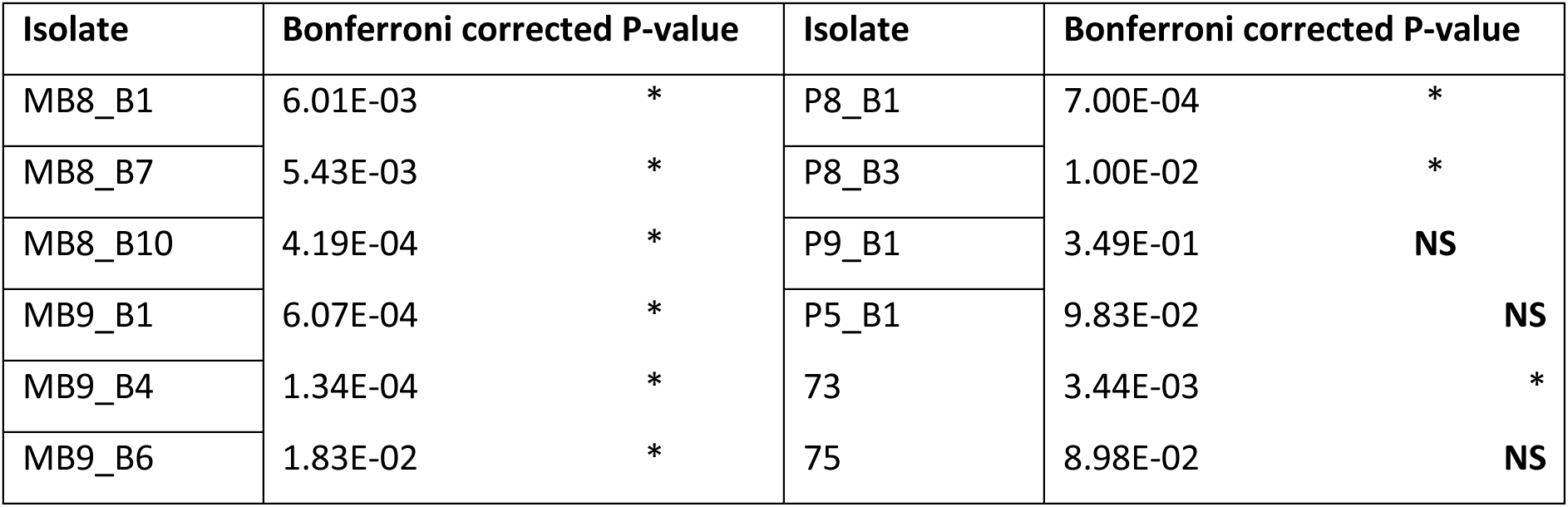
Bonferroni adjusted P-values from pairwise t-tests between pellicle timing of Δ*srfAC* control and with added 20 µM final concentration of surfactin, * = below significance cut-off 0.05, NS = not significant.

**Fig. S1.**
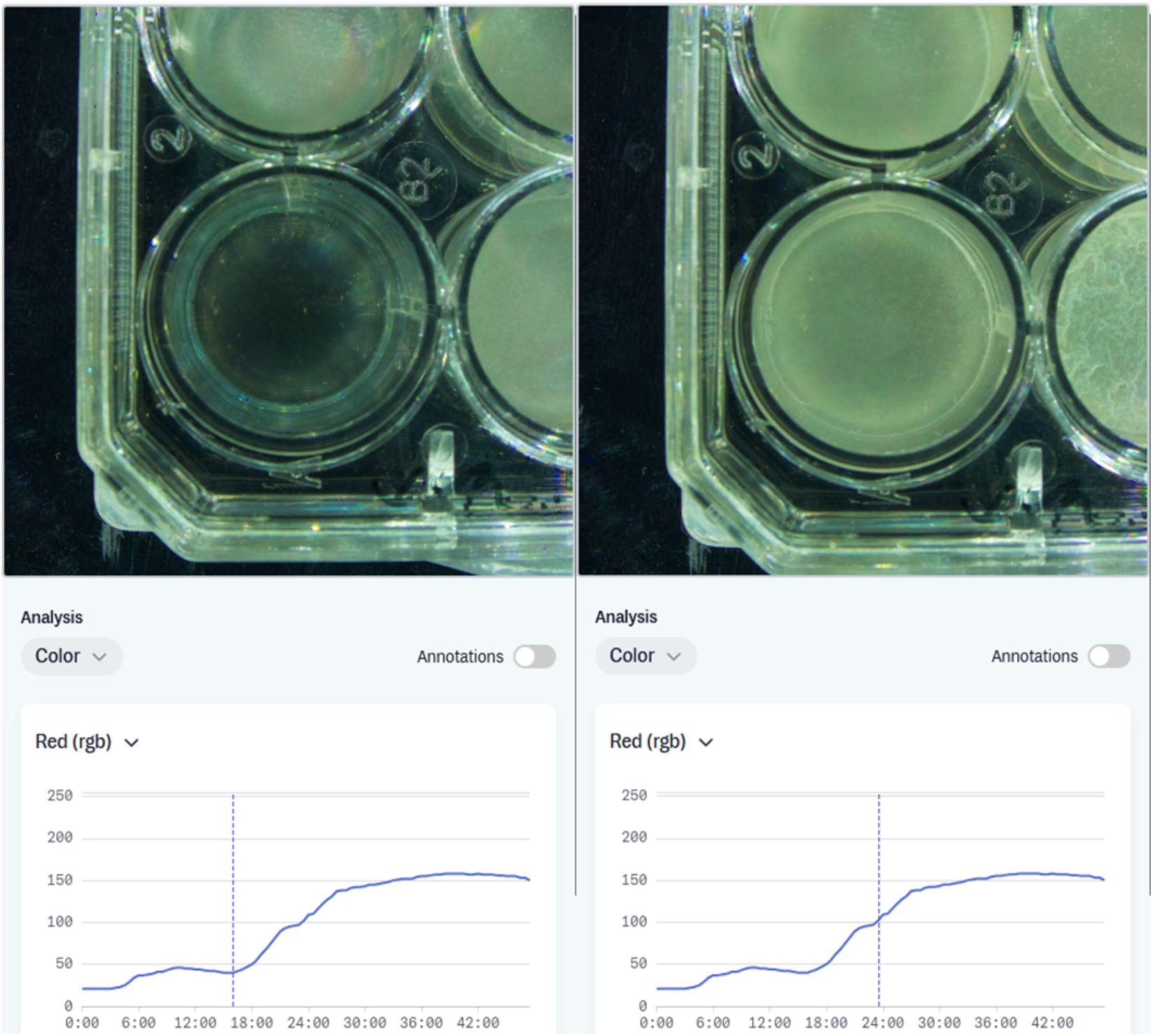
Screen capture showing pellicle culture of MB8_B1 with color analysis and red value before and at time of pellicle formation.

**Fig. S2.**
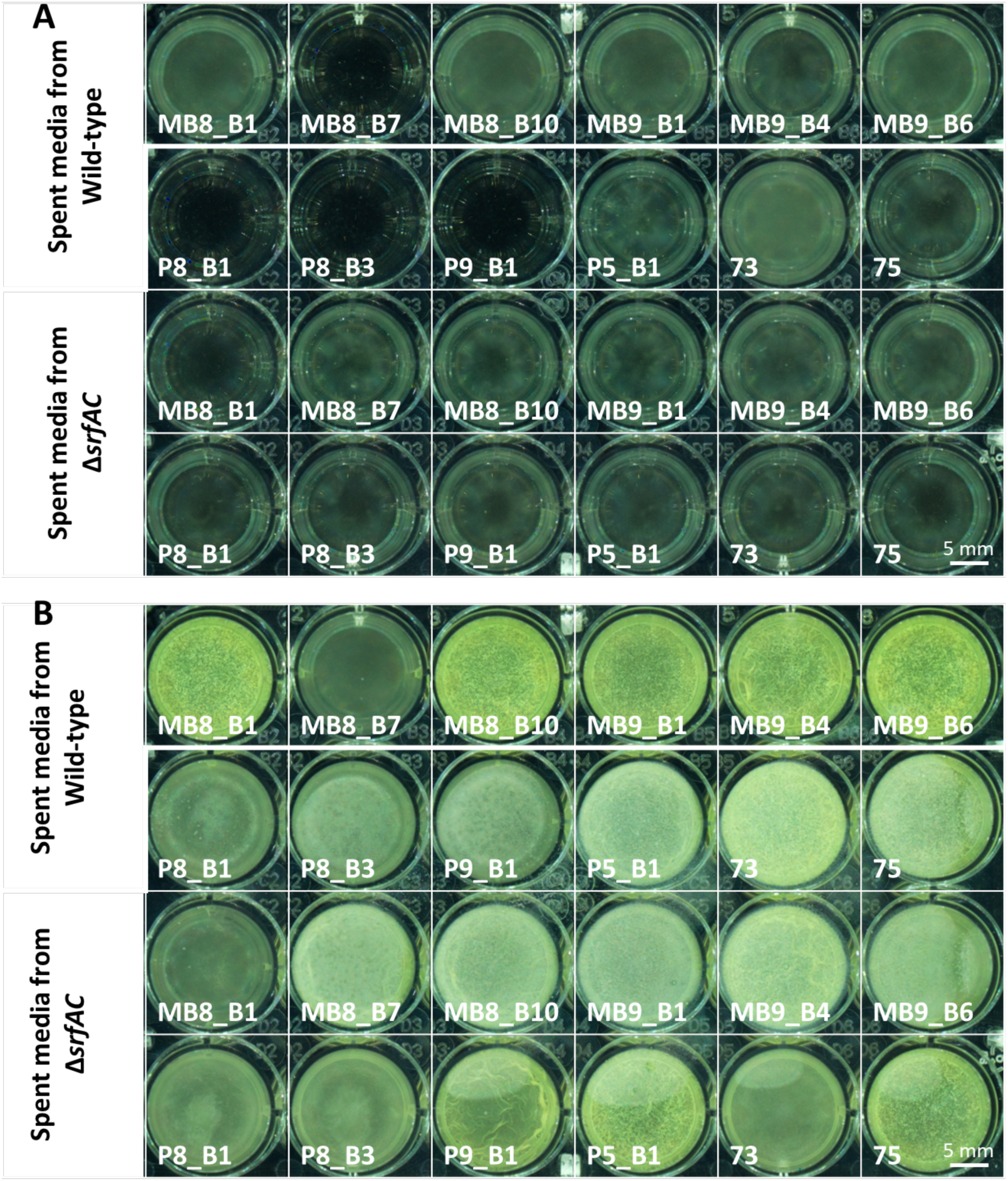
Inhibition by spent media from MB8_B7, P8_B1, P8_B3, and P9_B1 observed A) at 12 hours, B) and at 21 hours where pellicle is formed in spite of early inhibition.

**Fig. S3.**
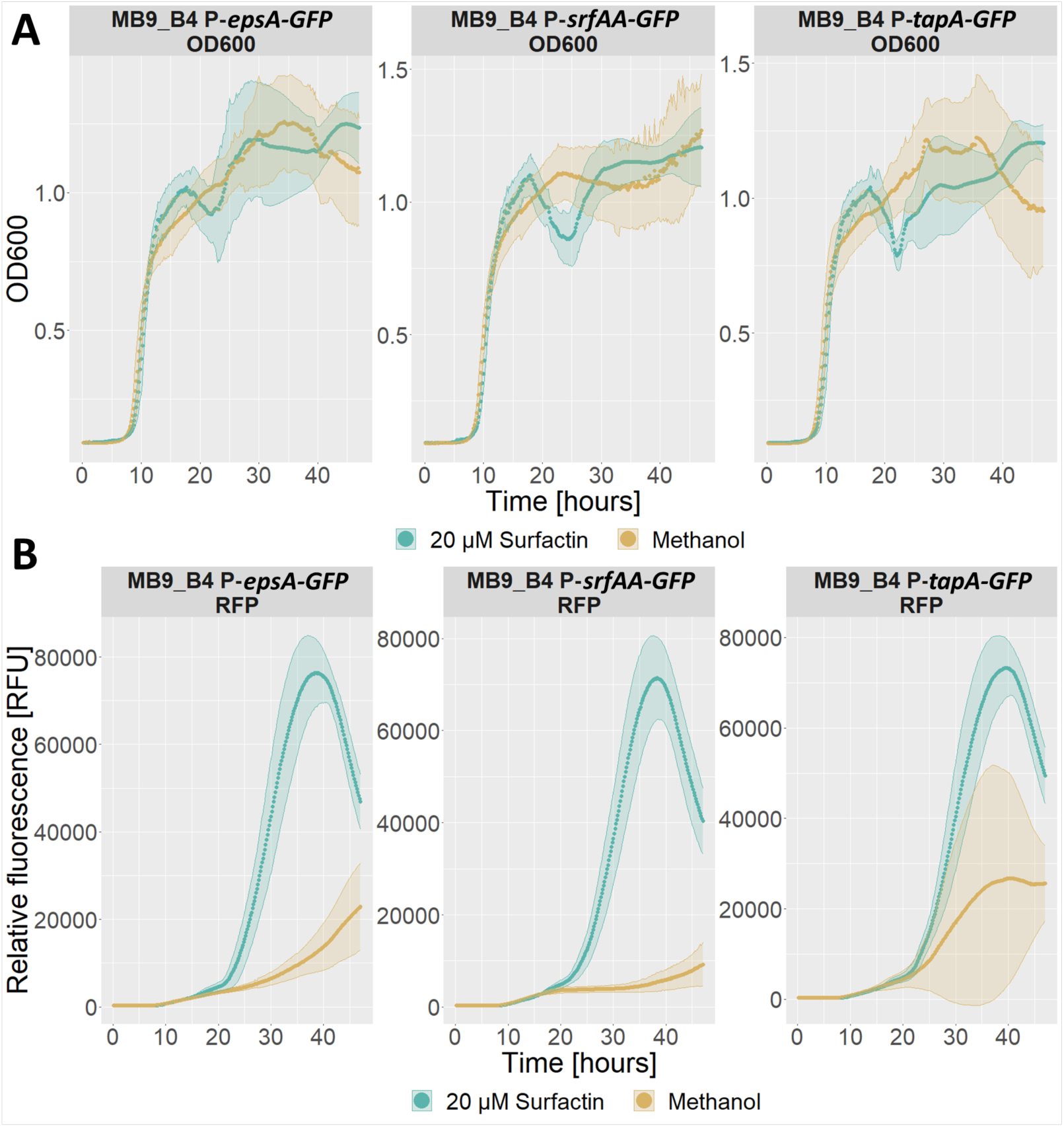
Effect of addition of surfactin to a final concentration of 20 µM (teal) or pure methanol as a control (beige) on growth and expression of fluorescence under constitutive and promoter-fusion coupled control in MB9_B4 *P-epsA-GFP8* (left), MB9_B4 *P-srfAA-GFP* (middle), and MB9_B4 *P-tapA-GFP* (right) over the course of 48 hours in LBgm at 30 °C. A) Optical density at 600 nm. B) Red fluorescence from RFP under control of a constitutive promoter. Data averaged from 3 biological replicates of each strain with ribbons showing standard deviation.

